# Interleukin-35 pretreatment attenuates lipopolysaccharide-induced heart injury by inhibition of inflammation, apoptosis and fibrosis

**DOI:** 10.1101/2020.01.23.916890

**Authors:** Yang Fu, Huan Hu, Meng Li, Huasong Xia, Yue Liu, Xiaopei Sun, Yang Hu, Fulin Song, Xiaoshu Cheng, Ping Li, Yanqing Wu

**Author notes:** Yang Fu and Huan Hu contributed equally. To whom correspondence should be addressed: Yanqing Wu: Department of Cardiology, The Second Affiliated Hospital of Nanchang University, No.1 Minde Road, Nanchang, 330006, China.

## Abstract

Previous studies have demonstrated that targeting inflammation is a promising strategy for treating lipopolysaccharide (LPS)-induced sepsis and related heart injury. Interleukin-35 (IL-35), which consists of two subunits, Epstein–Barr virus-induced gene 3 (EBI3) and p35, is an immunosuppressive cytokine of the IL-12 family and exhibits strong anti-inflammatory activity. However, the role of IL-35 in LPS-induced heart injury remains obscure. In this study, we explored the role of IL-35 in heart injury induced by LPS and its potential mechanisms. Mice were treated with a plasmid encoding IL-35 (pIL-35) and then injected intraperitoneally (ip) with LPS (10 mg/kg). Cardiac function was assessed by echocardiography 12 h later. LPS apparently decreased the expression of EBI3 and p35 and caused cardiac dysfunction and pathological changes, which were significantly improved by pIL-35 pretreatment. Moreover, pIL-35 pretreatment significantly decreased the levels of cardiac proinflammatory cytokines including TNF-α, IL-6, and IL-1β, and the NLRP3 inflammasome. Furthermore, increased BCL-2 levels and decreased BAX levels inhibited apoptosis, and LPS-induced upregulation of the expression of pro-fibrotic genes (MMP2 and MMP9) was inhibited. Further investigation indicated that pIL-35 pretreatment suppressed the activation of the cardiac NF-κBp65 and TGF-β1/Smad2/3 signaling pathways in LPS-treated mice. Similar cardioprotective effects of IL-35 pretreatment were observed in mouse myocardial fibroblasts challenged with LPS in vitro. In summary, IL-35 pretreatment can attenuate cardiac inflammation, apoptosis, and fibrosis induced by LPS, implicating IL-35 as a promising therapeutic target in sepsis-related cardiac injury.

---

Sepsis is a common medical emergency in hospital intensive care units. Over 30 million people worldwide are affected due to sepsis every year, representing a huge economic and public health burden in society (1). Sepsis-induced cardiomyopathy (SIC) is defined as heart injury and dysfunction occurring in patients with severe sepsis (2). Mechanistically, inflammation, oxidative stress, apoptosis and pyroptosis have been reported to serve as vital pathophysiology features of SIC (3, 4). Thus, finding potential therapeutic targets to selectively inhibit these pathological processes may help protect against sepsis and SIC.

Lipopolysaccharide (LPS), an important component of Gram-negative bacteria, can induce sepsis and related heart damage (5, 6). Inflammatory responses triggered by LPS, as evidenced by overproduction of proinflammatory cytokines (IL-6, IL-1β and TNF-α), are considered one of the major mechanisms underlying LPS-induced heart injury (7). Unresolved inflammation can lead to cardiac dysfunction, which has been reported to be related to higher rates of mortality among patients with sepsis in several clinical studies (8, 9). LPS also induces cardiac apoptosis by activating NOD-like receptor protein 3 (NLRP3) in the heart, and inhibition of apoptosis attenuates LPS-induced cardiac dysfunction (3, 10). Moreover, the adverse effects of LPS on cardiac fibroblasts (CFs), which account for two-thirds of all heart cells, have been reported to contribute to the progression of septic cardiac dysfunction (11, 12). LPS stimulation upregulates the expression of fibrosis signaling cascade proteins including metalloproteinases (MMPs) such as MMP2 and MMP9 in CFs to induce cardiac fibrosis, which exacerbates the cardiac dysfunction (13). Therefore, LPS may cause inflammatory responses, apoptosis, and fibrosis leading to heart injury, thus indicating that treatments of these three aspects of the pathological changes represents a therapeutic approach in LPS-induced heart injury.

Interleukin-35 (IL-35), which consists of two subunits, Epstein–Barr virus-induced gene 3 (EBI3) and IL-12A (also known as p35), is primarily secreted by regulatory T (Treg) and regulatory B (Breg) cells. This newly identified member of the heterodimeric IL-12 family has the capacity to suppress inflammation and immune responses (14, 15). IL-35 was reported to play a vital anti-inflammatory role in protecting against several inflammatory and autoimmune diseases including inflammatory bowel disease, chronic intestinal inflammation, rheumatoid arthritis, and asthma (16, 17, 18, 19). Mechanistically, IL-35 first interacts with its receptor (IL-35R), which is composed of three different dimers of IL-12Rβ2 and GP130, namely GP130-IL-12Rβ2, GP130-GP130, or IL-12Rβ2-IL-12Rβ2. This interaction induces signals within target cells that activate Janus kinase family members and phosphorylate members of this signal transduction pathway, which initiates the transcription of target genes to inhibit inflammatory responses (20, 21). To date, IL-35 has been reported to participate in preventing myocardial infarction (22), Coxsackie virus-B3-induced viral myocarditis (23), and doxorubicin and angiotensin II-induced cardiac injury (24, 25), implicating IL-35 as a promising therapeutic target for cardiovascular disease. However, the role of IL-35 in LPS-induced heart injury still remained obscure.

Our current study is the first to evaluate the effects of IL-35 pretreatment on inflammation, apoptosis, and fibrosis in heart injury induced by LPS in vivo and vitro.

## Results

### IL-35 expression decreased after LPS stimulation and increased after pIL-35 pretreatment

The results of Western blotting and qRT-PCR analyses confirmed expression of the IL-35 subunits (IL-12A/p-35 and EBI3) in mouse heart tissues after LPS stimulation and pIL-35 pretreatment. The levels of IL-12A/p35 and EBI3 protein expression in pIL-35(LPS-) mice were significantly (P < 0.05) elevated after pIL35 injection compared to those in the control mice (Figure 1A), demonstrating the ability of pIL-35 to upregulate IL-35 protein expression in mouse heart tissues. Further experiments revealed that the expression of EBI3 and IL-12A/p35 was significantly (P < 0.05) decreased at the protein and mRNA levels in mice after LPS stimulation, and the effect was reversed by pIL-35 pretreatment (Figure 1B-E).

**Figure 1.**
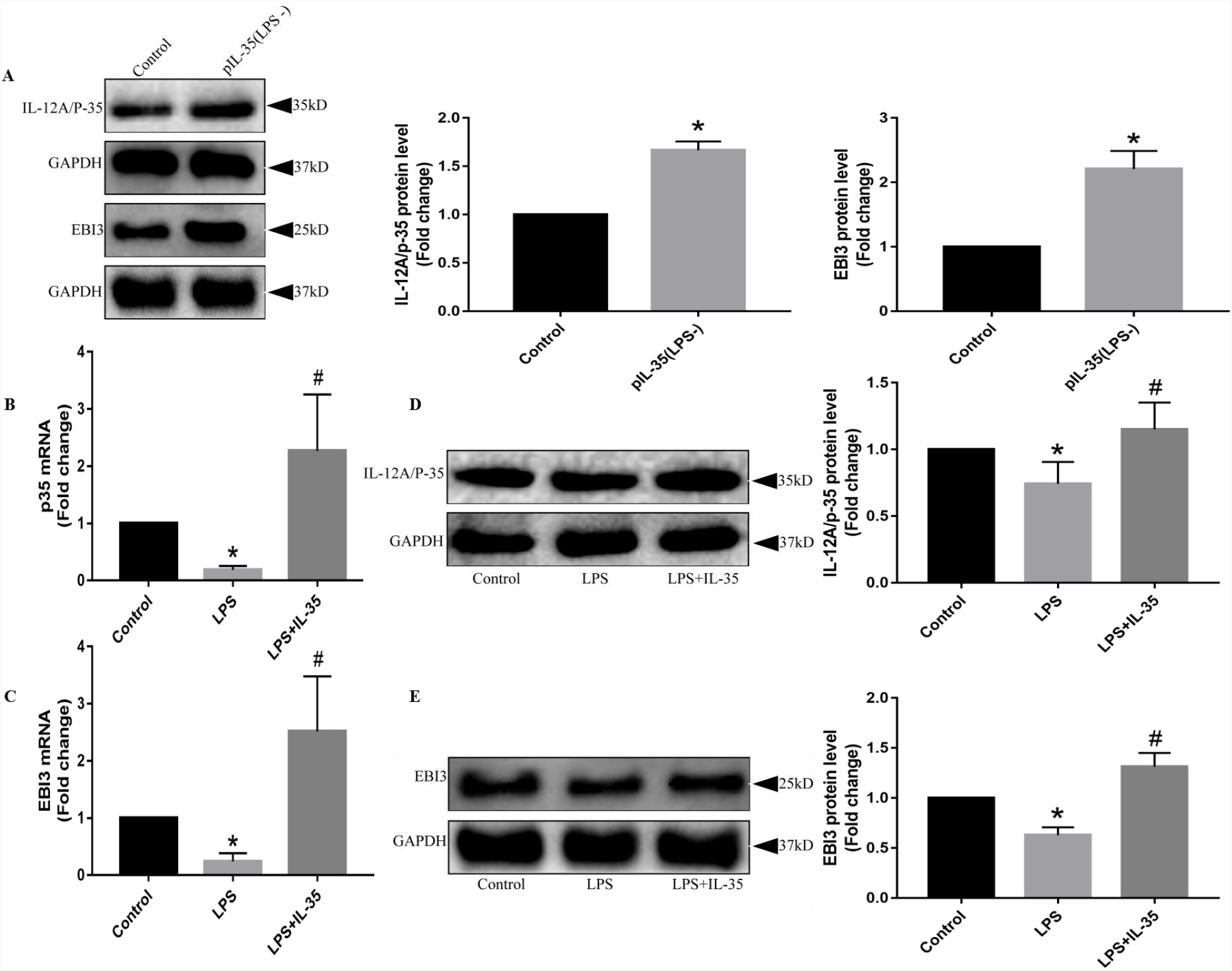
IL-35 levels were decreased in mouse heart tissue after LPS stimulation and increased after pIL-35 pretreatment. A, Western blot analysis of EBI3 and p35 protein expression levels in mouse heart tissue 7 days after pIL-35 pretreatment (n = 7–8). B and C, qRT-PCR analysis of EBI3 and p35 mRNA expression levels in mouse heart tissue after pIL-35 pretreatment and LPS stimulation (n = 7–8). D and E, EBI3 and p35 protein expressions in mouse heart tissue after pIL-35 pretreatment and LPS stimulation (n = 6-8). \**P* < 0.05 *vs.* the control group; #*P* < 0.05 *vs.* the LPS group. Data represent the mean ± SD of at least three experiments. Data were analyzed by one-way ANOVA and Student’s *t*-test.

### IL-35 pretreatment alleviated LPS-induced cardiac damage and dysfunction

The animal treatment protocols are summarized in Figure 2A. Mice pre-treated with pIL-35 were injected with LPS. The plasma levels of specific cardiac injury biomarkers (CK-MB and LDH) and histopathological changes were detected to evaluate the injury, and cardiac function was evaluated by echocardiography. Compared to the control group, the plasma CK-MB and LDH levels among the LPS-treated mice were increased, but were significantly (P < 0.05) decreased by the pIL-35 pretreatment (Figure 2B and C). Moreover, cardiac dysfunction, as evidenced by the decreased in the ejection fraction (EF%) and fractional shortening (FS%), was observed in the LPS-treated mice compared to that in the control mice, while the effects were significantly improved by pIL-35 pretreatment (Figure 2Da, E, and F). Similarly, compared to the control group, H&E and Masson’s trichrome staining revealed significant edema and disordered myofilament arrangement in myocardial cells and increased fibrosis in the heart tissues in the LPS group, while these pathological changes were significantly (P < 0.05) improved by pIL-35 pretreatment (Figure 2Db, Dc, and G). All these data suggested that IL-35 protects against LPS-induced cardiac damage and dysfunction.

**Figure 2.**
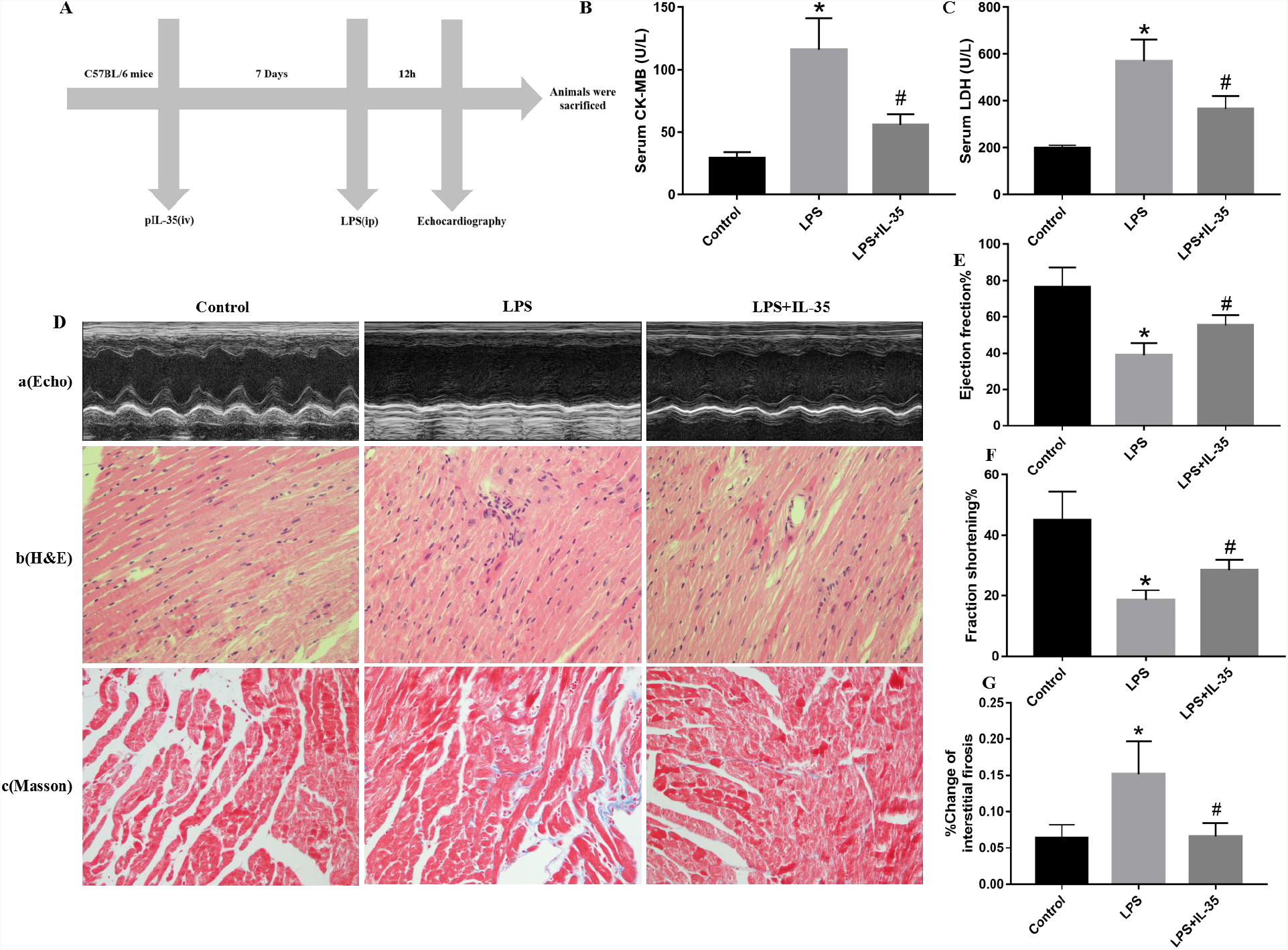
IL-35 pretreatment improved cardiac injury and function in mice stimulated with LPS. A, Animal treatment protocols used in the present study. B and C, Mouse serum was collected after LPS stimulation, and the production of CK-MB and LDH was detected (n = 5–8). Da, E and F, Echocardiographic analysis of ejection fraction (EF%) and fractional shortening (FS%) at the indicated points after LPS stimulation (n = 6–8). Db, HE staining of paraffin-embedded sections of heart tissues. Dc and G, Assessment of cardiac fibrosis by Masson’s trichrome staining paraffin-embedded sections of heart tissues. Quantification of fibrotic areas (n = 5–8). Data represent the mean ± SD. \**P* < 0.05 *vs.* the control group; #*P* < 0.05 *vs.* the LPS group.

### IL-35 pretreatment decreased LPS-induced cardiac inflammation

Abnormal anti-inflammatory and proinflammatory responses contribute to LPS-induced heart injury. To elucidate the effects of pIL-35 pretreatment on LPS-induced inflammatory responses, the levels of proinflammatory cytokines (IL-6, IL-1β and TNF-α), anti-inflammatory cytokine (IL-10), and the NLRP3 inflammasome in mouse heart tissue and serum were evaluated by qRT-PCR, Western blot and ELISA analyses. Compared to the control group, serum levels of IL-6, IL-1β and TNF-α levels were increased in LPS-treated mice, while pIL-35 pretreatment differentially inhibited these effects (Figure 3A-C). Similarly, the mRNA expression of IL-6, IL-1β and TNF-α in heart tissues was significantly (P < 0.05) upregulated in LPS-treated mice compared to the levels in the control group, while pIL-35 pretreatment efficiently decreased the three indices (Figure 3D–F). Similar inhibitory effects of pIL-35 pretreatment on the protein levels of proinflammatory cytokines (IL-6, TNF-α and MCP-1) and the NLRP3 inflammasome were also observed (Figure 3G-K). However, pIL-35 had no significant impact on plasma levels and mRNA expression of IL-10 (Figure 3L-M). IHC and immunofluorescence analysis of MCP-1 further confirmed that pIL-35 pretreatment inhibited LPS-induced inflammatory responses in mouse heart tissues (Figure S1A–C). All these data suggested that IL-35 protects against LPS-induced heart injury by inhibiting the excessive production of proinflammatory cytokines.

**Figure 3.**
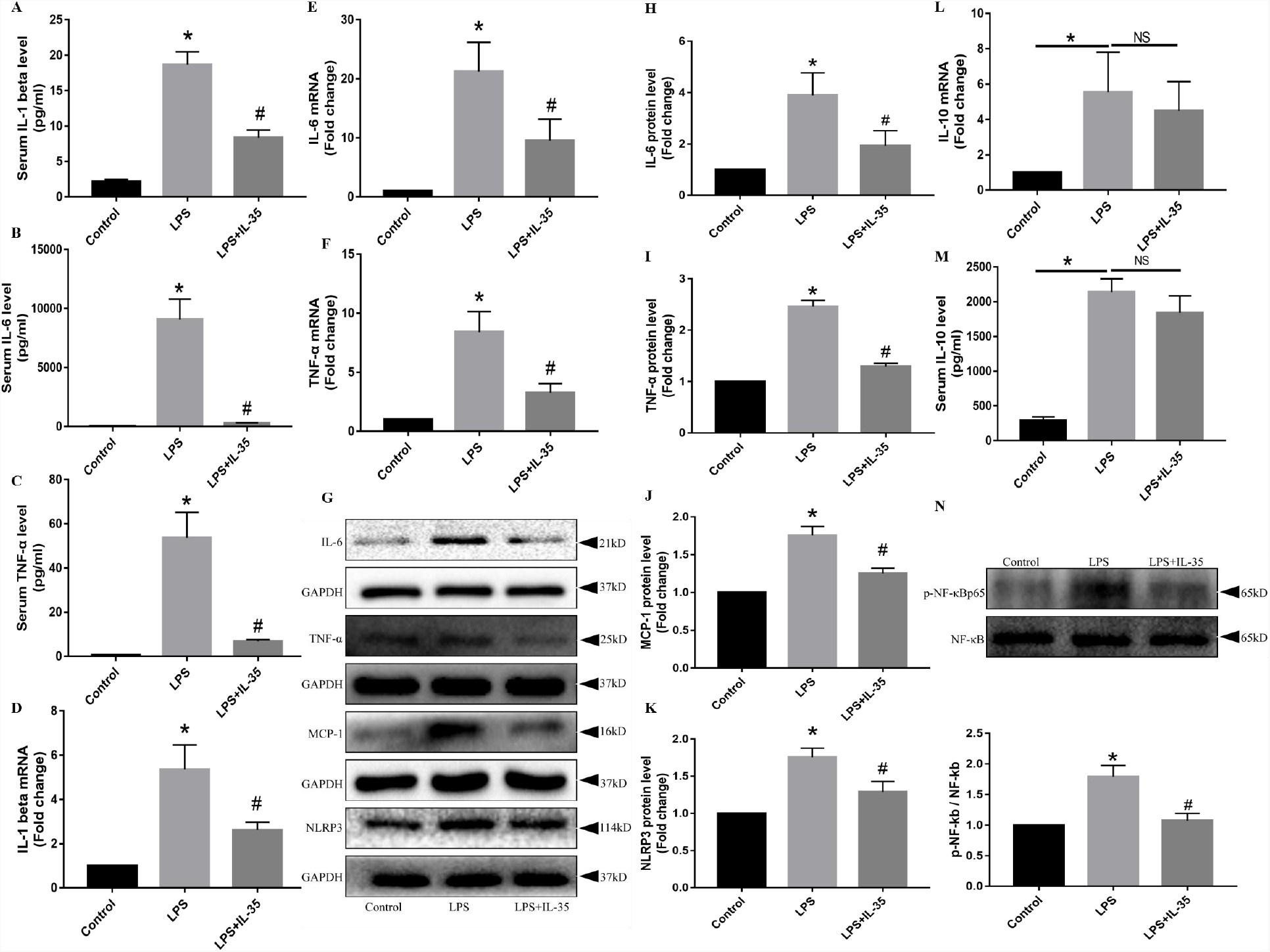
IL-35 pretreatment reduced the production of proinflammatory cytokines in the heart tissue and serum of mice stimulated with LPS. A–C, ELISA measurement of proinflammatory cytokines (IL-6, IL-1β and TNF-α) in each group (n = 5–8). D–F, qRT-PCR analysis of IL-6, IL-1β and TNF-α and cytokine IL-10 mRNA levels in mouse heart tissue. G-K, Western blot analysis of the proinflammatory cytokines (IL-6, TNF-α, MCP-1) and NLRP3 inflammasome protein levels in mouse heart tissue. L-M, Serum and heart tissues mRNA levels of anti-inflammatory cytokine IL-10 in each group. N, Protein expression levels of NF-κB/p65 and phosphorylation of NF-κB/p65 evaluated in mouse heart tissue. Data represent the mean ± SD of at least three experiments. \**P* < 0.05 *vs.* the control group; #*P* < 0.05 *vs.* the LPS group.

Proinflammatory cytokine production is known to be modulated by the NF-κB signaling pathway; thus, we further assessed the effect of pIL-35 pretreatment on the LPS-induced activation of the NF-κB signaling pathway in the mouse heart. As shown in Figure 3N, pIL-35 pretreatment suppressed the phosphorylation of the NF-κB/p65 component induced by LPS stimulation, indicating that IL-35 suppresses cardiac inflammatory responses partially by inhibiting activation of the NF-κB signaling pathway.

### IL-35 pretreatment inhibited apoptosis in the heart of LPS-treated mice

Overactive inflammatory responses usually lead to myocardial cell apoptosis. To investigate the effects of pIL-35 pretreatment on myocardial cell apoptosis in the mice with by LPS, we assessed the levels of the pro-apoptotic marker BAX and the anti-apoptotic marker BCL-2. Compared to the control mice, BAX expression was significantly (P < 0.05) increased at both the mRNA and protein levels in LPS-treated mice, and this effect was significantly (P < 0.05) inhibited by pIL-35 pretreatment (Figure 4A and B). In contrast, BCL-2 expression was significantly (P < 0.05) decreased at both the mRNA and protein levels in LPS-treated mice, while this effect was significantly (P < 0.05) improved by pIL-35 pretreatment (Figure 4C and D).

**Figure 4.**
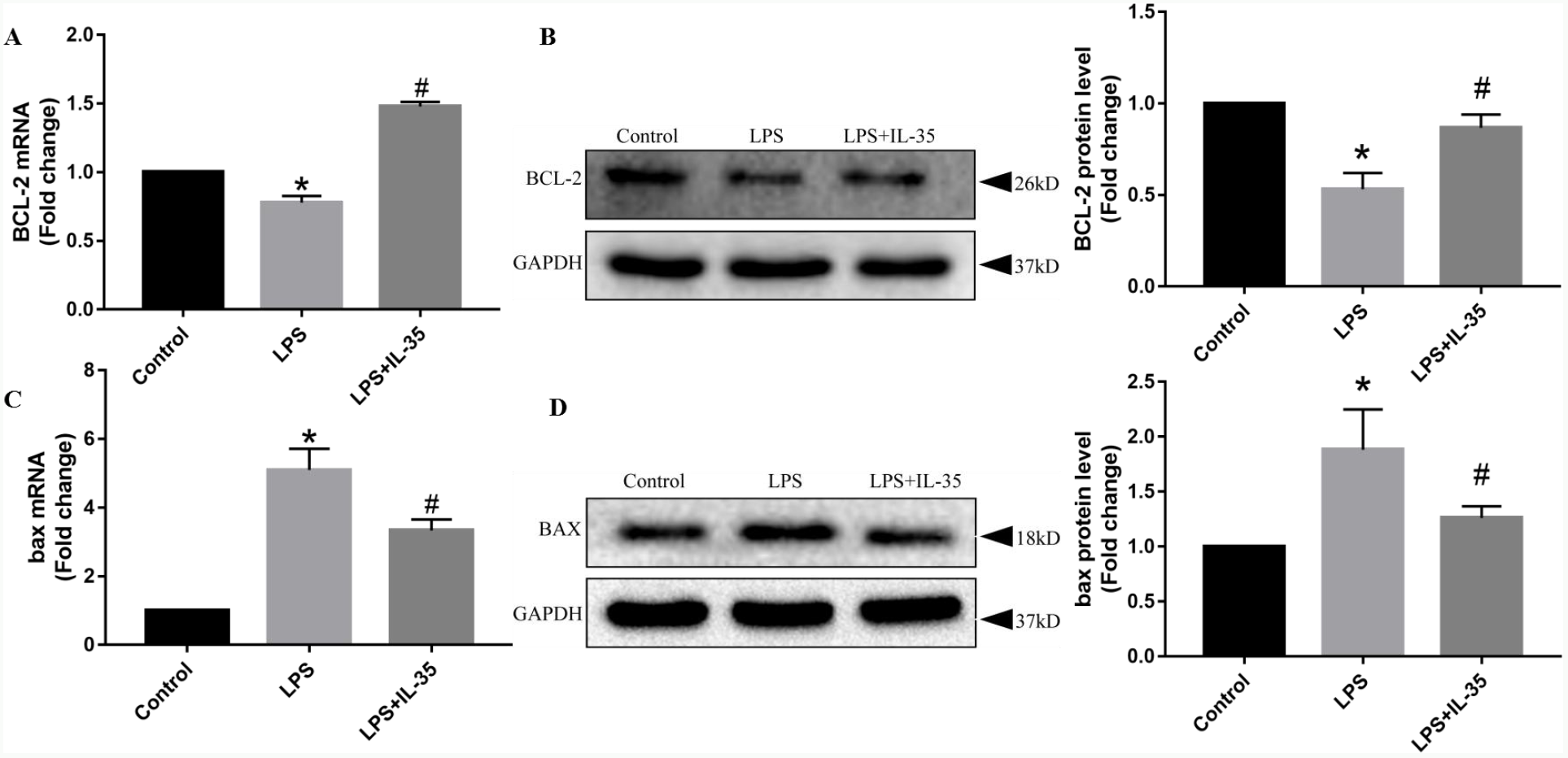
IL-35 pretreatment suppressed cardiac apoptosis in mice stimulated by LPS. A and B, Protein and mRNA levels of heart BCL-2 evaluated by Western blot and qRT-PCR analysis, respectively. C and D, Protein and mRNA expression levels of heart BAX assessed by Western blot and qRT-PCR analysis, respectively. Data represent the mean ± SD of at least three experiments. \**P* < 0.05 *vs.* the control group; #*P* < 0.05 *vs.* the LPS group.

### IL-35 pretreatment inhibited the LPS-induced cardiac fibrosis

Inflammatory responses and apoptosis are vital contributors to the development of cardiac fibrosis. To elucidate the effects of pIL-35 pretreatment of LPS-induced cardiac fibrosis, we evaluated the levels of fibrotic genes expressed in each group and observed that the expression of pro-fibrotic genes (MMP2 and MMP9) was significantly (P < 0.05) increased at both the mRNA and protein levels in the heart tissues of mice challenged with LPS compared to those in the control mice, while this effect was significantly (P < 0.05) inhibited by pIL-35 pretreatment (Figure 5A–D). Therefore, these data indicate that IL-35 exerts anti-fibrotic effects that protect against LPS-induced cardiac fibrosis.

**Figure 5.**
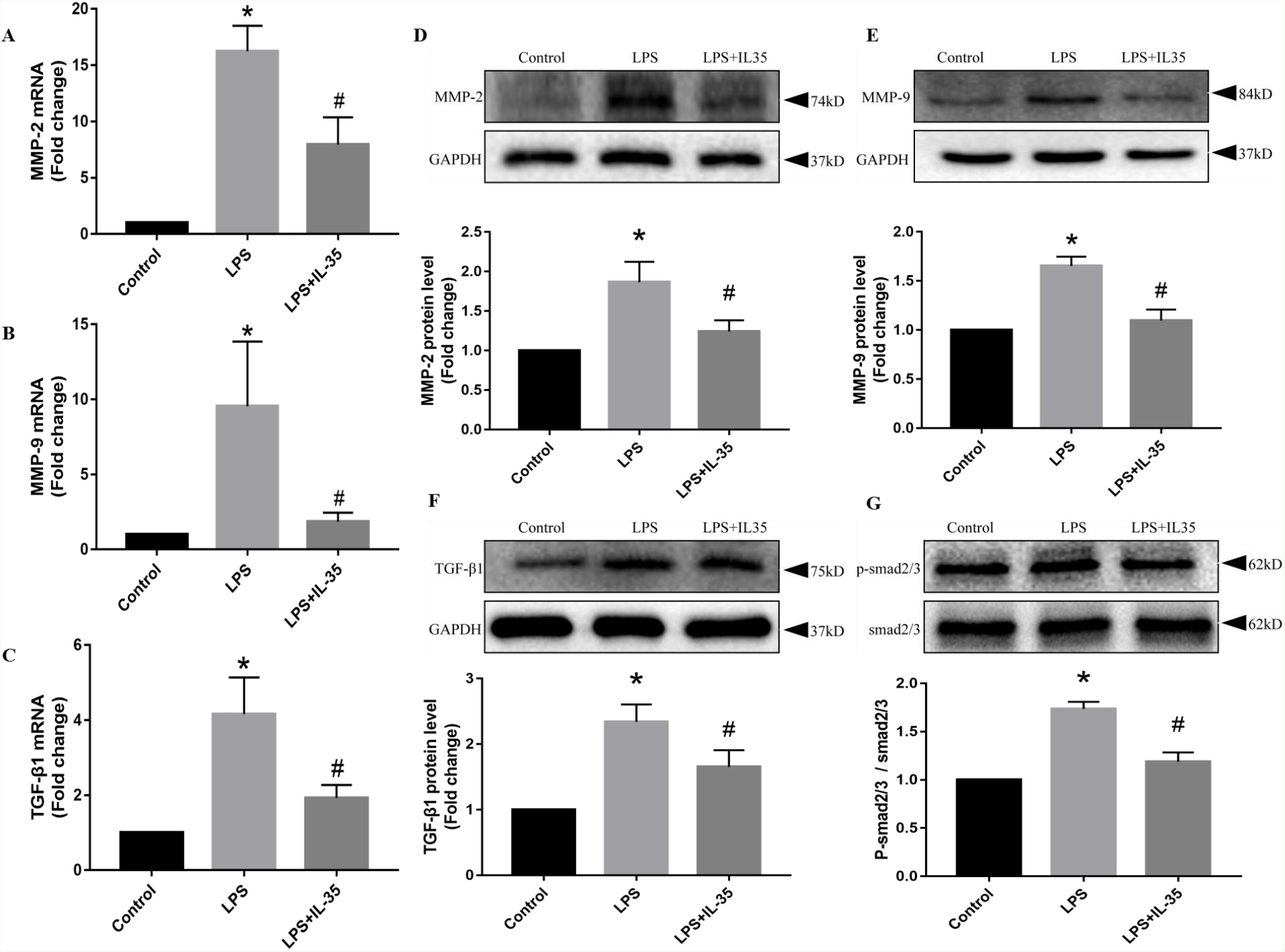
IL-35 pretreatment alleviated LPS-induced cardiac fibrosis in mice. A, B, D and E, Protein and mRNA levels of heart pro-fibrotic genes (MMP2 and MMP9) evaluated by Western blot and qRT-PCR analysis, respectively. C, F, and G, mRNA levels of heart TGF-β1 and protein levels of TGF-β1, total Smad2/3, and phosphorylation of Smad2/3 evaluated by Western blot and qRT-PCR analysis, respectively. Data represent the mean ± SD of at least three experiments. \**P* < 0.05 *vs.* the control group; #*P* < 0.05 *vs.* the LPS group.

The TGF-β/Smad signaling pathway plays a vital role in the process of fibrosis. Thus, the effects of pIL-35 pretreatment on the expression levels of components of this signaling pathway were further investigated by Western blot and qRT-PCR analyses. LPS stimulation upregulated the expression of TGF-β1 and the ratio of p-Smad2/3 to total Smad2/3, while the effect was inhibited by pIL-35 pretreatment (Figure 5E-G), indicating that the TGF-β1/Smad2/3 signaling pathway participates in the beneficial effects of pIL-35 pretreatment on LPS-induced cardiac fibrosis.

### IL-35 pretreatment reduced inflammation and fibrosis levels in cultured CFs stimulated by LPS in vitro

Cultured mouse CFs were pretreated with different concentrations of recombinant IL-35 protein (1 ng/ml, 10 ng/ml, and 50 ng/ml) for 1 h before adding LPS (100 ng/ml) for another 24 h. CCK8 assays of cell viability revealed significantly (P < 0.01) decreased viability in the LPS group compared with in the control group, while IL-35 pretreatment increased cell survival rate in a dose-dependent manner (Figure 6A). We further assessed whether IL-35 affected the cell survival rates by the inhibiting inflammatory response. Western blotting analysis revealed that IL-6, TNF-α and NLRP3 inflammasome proteins were significantly (P < 0.01) activated with increased expression levels in the LPS-treated cells compared to the control group, while IL-35 pretreatment differentially suppressed these indices (Figure 6B–E). Similar alterations in the expression of IL-6, IL-1β and TNF-α at the mRNA level were also observed (Figure S2A–C). Furthermore, IL-35 also inhibited the LPS-induced activation of the NF-κB/p65 signal pathway (Figure 6F).

**Figure 6.**
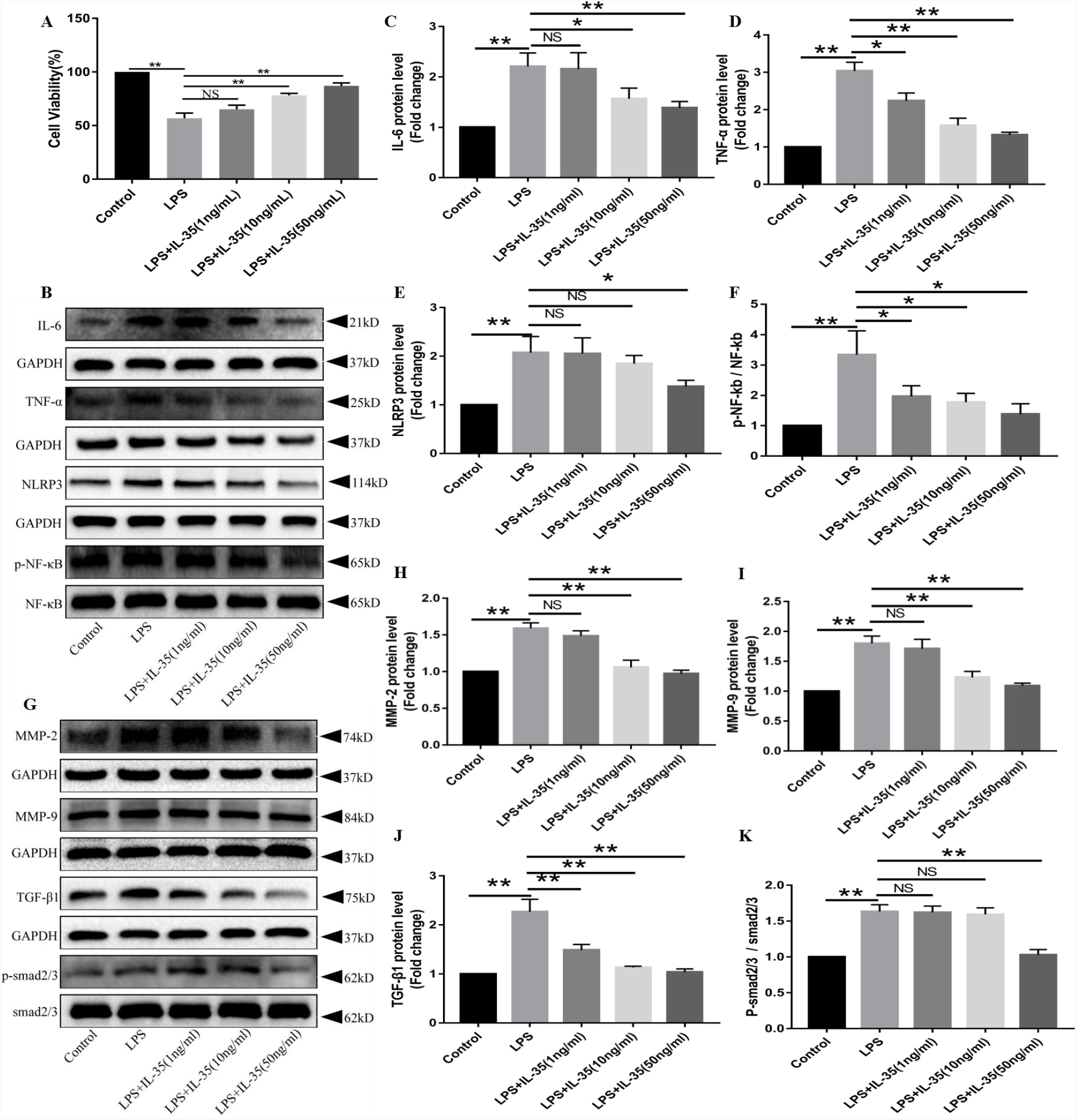
IL-35 pretreatment reduced inflammatory responses and fibrosis levels in LPS-treated CFs in vitro. A, CFs were incubated with IL-35 at three concentrations for 1 h before stimulation with LPS (100 ng/ml) for 24 h. Cell viability was then evaluated in CCK8 assays. B–F, Western blot analysis of protein levels of proinflammatory cytokines (IL-6 and TNF-α), the NLRP3 inflammasome, NF-κBp65 and phosphorylation of NF-κBp65. G–K, Western blot analysis of the protein levels of pro-fibrotic genes (MMP2 and MMP9), TGF-β1, total Smad2/3, and phosphorylation of Smad2/3. Data represent the mean ± SD of at least three experiments. \**P* < 0.05, \*\**P* < 0.01.

We also investigated the influence of IL-35 on the degree of fibrosis induced by LPS in CFs, a major cell type responsible for cardiac fibrosis. The protein expression of pro-fibrotic genes (MMP2 and MMP9) was significantly (P < 0.01) increased in the LPS group compared to the that in the control group, and this effect was reversed by IL-35 pretreatment (Figure 6G–I). Moreover, IL-35 pretreatment suppressed the LPS-induced expression of activated TGF-β1/Smad2/3 signaling pathway-related proteins (Figure 6G, J and K). Similar inhibitory effects of IL-35 on mRNA expression of MMP2, MMP9 and TGF-β1 were also observed (Figure S2D–F). Thus, these data indicated that IL-35 pretreatment decreases the degree of inflammation and fibrosis in LPS-treated CFs in vitro.

## Discussion

In the current study, we showed that IL-35 pretreatment protected against LPS-induced inflammatory responses, apoptosis, and fibrosis in the myocardium, implicating IL-35 as a promising therapeutic target in sepsis-induced cardiac injury.

IL-35 has been reported to have several important functions, such as anti-inflammatory, anti-oxidative, and anti-apoptotic properties. IL-35 protected against acute liver and kidney injury caused by LPS through inhibiting inflammatory responses (27, 28). IL-35 also inhibited lysophosphatidylcholine-induced mitochondrial reactive oxygen species (mtROS) production and human aortic endothelial activation (29, 30). Furthermore, IL-35 was reported to exert anti-apoptotic effects that prevent diabetic neuropathic pain and doxorubicin-induced cardiac injury (24, 31). Recently, Jia, et al.(22) found that IL-35 reduced cardiac rupture, improved wound healing, and attenuated cardiac remodeling after myocardial infarction by promoting reparative CX3CR1^+^Ly6C^low^ macrophage survival. Moreover, Li et al.(25) found that interleukin-12p35 deletion facilitated CD4 T-cell-dependent macrophage differentiation and enhanced angiotensin II-induced cardiac fibrosis. Therefore, we speculated that IL-35 might exert cardioprotective effects in sepsis. Thus, in this study, we established in vitro and in vivo models of LPS-induced cardiac damage to demonstrate the potential cardioprotective properties of IL-35. We found that IL-35 pretreatment alleviated the injury to the heart induced by LPS, including cardiac dysfunction and myocardial pathological changes, through inhibiting proinflammatory responses, apoptosis, and fibrosis. Furthermore, our results indicated the involvement of the NF-κB and TGF-β1/Smad2/3 signaling pathways in the cardioprotective effects of IL-35.

IL-35 pretreatment suppressed LPS-induced inflammatory responses in heart tissues and cultured CFs. Although the molecular mechanisms of SIC remain to be fully elucidated, inflammatory responses are acknowledged as a main contributor to the occurrence and development of SIC (32, 33). In this study, LPS acted as a vital contributor to NF-κB activation, which can lead to the transcription of proinflammatory cytokines such as IL-6, IL-1β and TNF-α (34). Several previous studies have indicated that IL-35 serves as an anti-inflammatory factor in anti-atherosclerotic therapy and inhibited LPS-induced vascular endothelial cell activation and inflammation (35, 36), suggesting that IL-35 might prevent SIC by suppressing inflammation. The present study is the first to show that IL-35 pretreatment significantly attenuated LPS-induced heart proinflammatory responses, as evidenced by the decreased production of proinflammatory cytokines in the serum, heart tissues and cultured CFs. The effects of IL-35 on production of the anti-inflammatory cytokine IL-10 are controversial (23, 27, 28). In this study, we did not observe any effects of IL-35 on the levels of the anti-inflammatory cytokine IL-10 in mouse heart tissue and serum (Figure 3L, and M), suggesting that the anti-inflammatory properties of IL-35 are not dependent on IL-10. Thus, our findings indicate that IL-35 possesses the capacity to protect against LPS-induced heart injury by inhibiting inflammation.

Apoptosis is reported to be involved in SIC. Previous studies have indicated that LPS-induced inflammatory responses cause apoptosis (37, 38). Liu, et al.(39) found that treatment by Naringin decreased the ratio of BAX to BCL-2 to withstand apoptosis and prevent LPS-induced cardiac injury. Actually, the balance of pro-apoptotic and anti-apoptotic proteins plays a vital role in LPS-induced cardiac apoptosis (40). Expression of the pro-apoptotic family protein BAX is increased by LPS stimulation, while the restraint of BAX expression protects against LPS-induced heart injury (41). In contrast, the anti-apoptotic family protein BCL-2 inhibits apoptosis to protect against LSP-induced heart injury. Zhang et al. (42) demonstrated that BCL-2 levels were decreased in heart tissues following LPS stimulation, while Apigenin increased BCL-2 protein expression to alleviate cardiac damage. Therefore, it can be speculated that an imbalance in the levels of BAX and BCL-2 proteins contributes to LPS-induced cardiac damages, and treatments that improve this imbalance might protect against LPS-induced cardiac damage. Similarly, increased apoptosis, as evidenced by the increase in BAX levels and decrease in BCL2 levels, was also observed in the heart injury stimulated by LPS in this study, while pIL-35 pretreatment significantly reversed the effects. Thus, the LPS-induced cardiac damage was shown to be associated with the imbalance between BAX and BCL-2, indicating that the cardioprotective effects of IL-35 are also relevant to it is anti-apoptotic properties.

Myocardial fibrosis contributes to the development of LPS-induced cardiac dysfunction (43). CFs plays pivotal roles in regulating both the normal physiology and the pathological condition of heart tissues(44). To date, numerous studies have implicated IL-35 as a new anti-fibrotic effector in liver fibrosis, skin fibrosis, pulmonary fibrosis and renal fibrosis, while the effects of IL-35 in cardiac fibrosis remain unclear (45). In this study, we showed that the expression of several pro-fibrotic genes was increased in LPS-induced cardiac injury, which is consistent with previous reports (46). Our findings further confirmed the ability of IL-35 to attenuate LPS-induced cardiac fibrosis in mice and cultured CFs, as evidenced by the decreased cardiac fibrosis area and levels of pro-fibrotic genes (MMP2 and MMP9). The TGF-β/Smad signaling pathway has been reported to exert a well-known effect on the development of cardiac fibrosis. Since previous research indicated that IL-35 serves as a TGF-β signal antagonist in liver fibrosis, and cardiac fibrosis has been proved to share many common features with liver fibrosis(47, 48), we speculated that TGF-β/Smad signaling might be involved in the anti-fibrotic effects of IL-35 in LPS-induced heart fibrosis. Our findings provide the first evidence that IL-35 pretreatment suppresses activation of the TGF-β1/Smad2/3 signaling pathway during the progression of LPS-induced cardiac fibrosis, as evidenced by the decreased expression of TGF-β1 and ratio of p-Smad2/3 to total Smad2/3.

In conclusion, this is the first study to demonstrate that IL-35 pretreatment prevents sepsis-induced cardiac injury by inhibiting inflammatory responses, apoptosis, and cardiac fibrosis. Moreover, our results indicate that the NF-κBp65 and TGF-β1/Smad2/3 signaling pathways are involved in the cardioprotective effects of IL-35. Therefore, the findings of this study indicate that IL-35 is a promising target to protect against cardiac injury in sepsis.

## EXPERIMENTAL PROCEDURES

### Materials

In the present study, we used a total of 40 male C57BL/6 J mice (aged 8–10 weeks, weighing 22 ± 3g) obtained from Hunan SJA Laboratory Animal Co., Ltd., Changsha, China. Plasmid pcDNA3.1-IL-35 (pIL-35) (Shanghai Genechem Co., Ltd., Shanghai, China) was constructed and used for the treatment of mice. Recombinant IL-35 cytokine (Cat# 8608-IL, R&D Systems, Minnesota, USA) was obtained and used for cultured CFs treatment. The following primary antibodies used in the present study: anti-EBI3 (Cat# ab124694), anti-IL-12A/P35 (Cat# ab131039), anti-IL-6 (Cat# ab6672), anti-TNF-α (Cat# ab1793), anti-MCP-1 (Cat# 66272-1-Ig), anti-NLRP3 (Cat# 19771-1-AP), anti-NF-κBp65 (Cat# CST-8242S), anti-p-NF-κBp65 (Cat# CST-3033S), anti-BCL2 (Cat# 12789-1-AP), anti-BAX (Cat# 50599-2-Ig), anti-TGF-β1 (Cat# 21898-1-AP), anti-Smad2/3 (Cat# CST-5678S), anti-p-Smad2/3 (Cat# CST-9510S), anti-MMP2 (Cat# ab97779), anti-MMP9 (Cat# ab58803), and anti-GAPDH (Cat# 60004-1-Ig). Plasma lactate dehydrogenase (LDH) and creatine kinase-MB (CK-MB) activity in mice were measured using an automatic biochemical analyzer (Chemray240, Rayto Life Technology Co., Ltd., Shenzhen, China). The enzyme-linked immunosorbent assay (ELISA) kits for detection of IL-1β (Cat# 88-7013), IL-6 (Cat# 88-7064), and TNF-α (Cat# 88-7324) were obtained from Invitrogen, California, USA, and the kit for detection of IL-10 (Cat# EK210/3-01) was purchased from MultiSciences Biotech Co., Ltd., Wuhan, China. Lipopolysaccharide (LPS) was obtained from Sigma (St. Louis, MO, USA).

### Animal treatment

All animal treatments were performed in accordance with the institutional guidelines of the Animal Care and Use Committee of the Second Affiliated Hospital of Nanchang University (China). Animals were maintained 23 ± 2°C and a relative humidity of 60 ± 10% under a 12-h light/dark cycle, and with free access to water and rodent chow.

After adaptation to the housed environment for 1 week, mice were then randomized into two groups: (1) Control group (Control, n = 8): phosphate buffered saline (PBS) was injected via the tail vein; (2) IL-35 group {IL35(LPS-), n = 8}: pIL-35 was injected via the tail vein and without LPS treatment to confirm the IL-35 overexpression in heart tissues. The remaining mice were randomized into another three groups: (1) Control group (Control, n = 8): this control group was pretreated with PBS and with no further treatment; (2) LPS group (LPS, n = 8): the mice pretreated with PBS were injected with LPS (10 mg/kg, ip); (3) LPS + IL-35 group (LPS+IL-35, n = 8): the mice pretreated with pIL-35 were injected with LPS (10 mg/kg, ip). According to previous research,^26^ a hydrodynamic-based gene transfection technique involving rapid tail vein injection of 100 μg plasmid DNA in 2.0 ml PBS was used to treat mice. Furthermore, the protocols used to treat the housed animals are summarized in Figure 2A.

### Echocardiography

Echocardiography was conducted to evaluate the changed in cardiac function in each group. First, mice were anesthetized by continuous administration of isoflurane at 1.5%, fastened to a heating pad in the supine position and the chest hair was shaved. Transthoracic echocardiography was then conducted using a Vevo770 (VisualSonics, Toronto, Canada) carrying a 30 Hz transducer. We used the short-axis view under M-mode tracings to detect left ventricle (LV) dimensions during the course of systole and diastole. Subsequently, we measured LV end systolic and diastolic diameter (LVEDS, LVEDD), LV end systolic and diastolic posterior wall thickness (LVPWS, LVPWDD). According to the LVEDS and LVEDD data, the ejection fraction (EF%) and shortening fraction (FS%) were gained via automatic calculation.

### Culture of mouse CFs

Mouse CFs cells were seeded in 6-cm-diameter dishes at a density of 1×10^6^cells and cultured in Dulbecco’s modified Eagle’s medium (DMEM) supplemented with 10% fetal bovine serum (FBS) and 1% penicillin/streptomycin (Hyclone) in a humidified incubator at 37°C under 5% CO_2_. Before any experimental treatment, the cells should be grown to 90% confluence. Available CFs were pretreated with IL-35 at low (1 ng/ml), medium (10 ng/ml), and high (50 ng/ml) concentration for 1 h and then challenged with LPS (100 ng/ml) for another 24 h. In this study, available CFs were randomized into five groups: (1) Control group: no treatment; (2) LPS group: LPS (100 ng/ml); (3) Low-dose IL-35 pretreatment+LPS group: LPS+IL-35 (1 ng/ml), with the addition of IL-35 (1 ng/ml) and LPS (100 ng/ml); (4) Medium-dose IL35 pretreatment+LPS group: LPS+IL-35 (10 ng/ml), with the addition of IL-35 (10 ng/ml) and LPS (100 ng/ml); (5) High-dose IL35 pretreatment+LPS group: LPS+IL-35 (50 ng/ml), with the addition of IL-35 (50 ng/ml) and LPS (100 ng/ml). After treatment, the cells in each group were harvested for further molecular investigation.

### CCK8 assay

The CFs (2 × 10^5^ cells/ml) pretreated with different concentrations of IL-35 were seeded in 96-well plate supplemented with 100 μl DMEM each well. Then the CFs were challenged with LPS (100 ng/ml) for 24h. Subsequently, cell medium of each well was replaced with 100 μl fresh medium, which contained 10 μl CCK8 reagent (Cat# C0038, Beyotime Biotechnology, Shanghai, China), and further incubated for 2-3 h. Finally, a microplate reader (Thermo Fisher Scientific, Massachusetts, USA) was used to measure the absorbance of each well, and cell viability of each group was calculated according to the standard method. This experiment was conducted more than three times in independent experiments.

### LDH and CK-MB activity assays

After anesthetization with pentobarbital sodium (30mg/kg, ip), mice were sacrificed and whole blood was obtained from the right ventricle with vacuum tubes. Subsequently, whole blood was centrifuged at 3,000 rpm for 20 min at 4°C to collect the plasma. The plasma levels of the cardiac injury markers CK-MB and LDH were measured using an automatic biochemical analyzer (Chemray240, Rayto, China).

### Enzyme-linked immunosorbent assay (ELISA)

Mouse plasma levels of proinflammatory cytokines (IL-1β, TNF-α, and IL-6), and the anti-inflammatory cytokine IL-10 were detected using specific mouse IL-1β (Invitrogen, California, USA), IL-6 (Invitrogen, California, USA), TNF-α (Invitrogen, California, USA), and IL-10 (MultiSciences Biotech Co., Ltd., Wuhan, China) ELISA kits according to the standard procedures.

### Histopathological assessment of the heart tissues

Mouse heart tissue was excised immediately after sacrifice, the aorta and atria were separated carefully, and the tissue was washed three times in ice-cold saline. One portion of the heart was fixed in 4% formaldehyde solution for 1 h and cut into 5-μm sections at various depths to prepare for staining with hematoxylin and eosin (H&E), Masson’s trichrome (Masson), as well as immunofluorescence and immunohistochemistry (IHC) analyses according to standard procedures. The remaining heart tissues were stored at −80°C for further molecular analysis.

### RNA extraction and qRT-PCR

Total RNA was extracted from heart tissues and cultured CFs using TRIzol reagent (Cat# DP424, Tiangen Biotech, China), and cDNA transcription was accomplished using a first strand cDNA synthesis kit (Cat# KR116-02, Tiangen Biotech, China). Subsequently, quantitative real-time PCR (qRT-PCR) was performed with SYBR green as fluorescence dye (Cat# FP205-02, Tiangen Biotech, China) in a total reaction volume of 20 μL according to the standard manufacturer’s instructions. Details data of the primers used in this study are shown in Table S1. The qRT-PCR analysis was conducted using a PCR thermocycler (ABI7900TH, USA) and relative gene expression was calculated using the 2^−ΔΔCT^method; GAPDH content was quantified as an internal control. All primer sequences were designed and constructed by Shanghai Sangon Biotech Engineering Co., Ltd., Shanghai, China.

### Western blot analysis

Total proteins were extracted from mouse heart tissues and cultured CFs using a specific protein extraction kit (Cat# P0013B, Beyotime Biotechnology, Jiangsu, China). Protein (60 μg per sample) were separated by 10% sodium dodecyl sulfate-polyacrylamide gel electrophoresis (SDS-PAGE) and then transferred to Polyvinylidene fluoride (PVDF) membranes (Millipore, Bedford, MA, USA). Membranes were then blocked for 90 min with specific 5% non-fat dried milk and incubated with primary detection antibodies, including anti-EBI3 (1:1000), anti-IL-12A/P35 (1:1000), anti-IL-6 (1:500), anti-TNF-α (1:500), anti-MCP-1 (1:1000), anti-NLRP3 (1:800), anti-NF-κBp65 (1:1000), anti-p-NF-κBp65 (1:500), anti-BCL2 (1:800), anti-BAX (1:1000), anti-TGF-β1 (1:1000), anti-Smad2/3 (1:1000), anti-p-Smad2/3 (1:500), anti-MMP2 (1:1000), anti-MMP9 (1:000), and anti-GAPDH (1:1000), overnight before incubation with specific mouse or rabbit peroxidase-labeled secondary antibodies (1:5000). Protein bands were then detected with an enhanced chemiluminescence (ECL) detection kit and an ECL detection instrument (Thermo Fisher Scientific). Image Lab 4.0.1 was used to analyze these results. In this study, all proteins measurements were repeated more than three times in independent experiments.

### Statistical analysis

All statistical analysis was conducted with GraphPad Prism 7.0 software (GraphPad Software Inc., San Diego, CA, U. S.). The data were represented as means ± standard deviation (SD). Student’s *t*-test was used to compare the results of two groups. Furthermore, one-way ANOVA was used to evaluate differences between two or more groups. In the present study, *P* values < 0.05 were considered to indicate statistical significance.

## Acknowledgments

The present study was supported by the National Key Research and Development Plan of China (2016YFC1301100), the National Natural Science Foundation of China (81660062, 81560079, 81860058), the Funding Scheme for Academic and Technical Leaders of Major Disciplines, Jiangxi Province, China (20113BCB22005, 20172BCB22027), Beijing Health Promotion Association (20181BCB24013), and special funds for guiding local scientific and technological development by the central government of China (S2019CXSFG0016).

## Compliance with ethical standards

Conflict of Interest. The authors declare that they have no conflict of interest.

